# ipcoal: An interactive Python package for simulating and analyzing genealogies and sequences on a species tree or network

**DOI:** 10.1101/2020.01.15.908236

**Authors:** Patrick F. McKenzie, Deren A. R. Eaton

## Abstract

**Summary:** *ipcoal* is a free and open source Python package for simulating and analyzing genealogies and sequences. It automates the task of describing complex demographic models (e.g., with divergence times, effective population sizes, migration events) to the *msprime* coalescent simulator by parsing a user-supplied species tree or network. Genealogies, sequences, and metadata are returned in tabular format allowing for easy downstream analyses. *ipcoal* includes phylogenetic inference tools to automate gene tree inference from simulated sequence data, and visualization tools for analyzing results and verifying model accuracy. The *ipcoal* package is a powerful tool for posterior predictive data analysis, for methods validation, and for teaching coalescent methods in an interactive and visual environment.

**Availability and implementation:** Source code is available from the GitHub repository (https://github.com/pmckenz1/ipcoal/) and is distributed for packaged installation with conda. Complete documentation and interactive notebooks prepared for teaching purposes are available at https://ipcoal.readthedocs.io/.

## 1 Introduction

The coalescent process (Hudson, 1983; Kingman, 1982) is used to model the distribution of genealogical ancestry across a set of sampled genomes. It approximates a neutral Wright-Fisher process of random mating within populations where the expected waiting times between subsequent coalescent events can be drawn from a statistical distribution based on the effective population size. This makes simulation of genealogies under the coalescent process (Hudson, 2002) a computationally efficient approach for integrating over genealogical variation (i.e., treating it as a latent random variable) when making population genetic inferences (Beerli & Felsenstein, 2001).

Demographic models specify the parameters of a coalescent simulation. Highly complex models may include population sizes and divergence times, and gene flow (admixture) between populations. For example, in the study of human history, a demographic model may describe divergences among different continents, the expansion of populations separately in Africa, Eurasia, and the Americas, and subsequent admixture between them (Reich, 2018; Gronau *et al.*, 2011; Green *et al.*, 2010). Demographic models are also routinely used in phylogenetics, with the goal of inferring a topology (i.e., the relationships among connected populations) in addition to the parameters of a demographic model applied to the topology (Knowles & Kubatko, 2011; Degnan & Rosenberg, 2009).

The ability to simulate realistic sequence data evolving on genealogies sampled from complex demographic models has enabled new types of inference from genomic data, from fitting parameters to demographic models and performing model comparisons (Chung & Hey, 2017); to performing posterior predictive data analyses (Brown, 2014); to generating training datasets for machine learning methods (Schrider & Kern, 2018); to validating new inference methods (Adrion *et al.*, 2019). Despite the impressive capabilities of recent state-of-the-art coalescent simulation tools like *msprime* (Kelleher *et al.*, 2016), it is difficult for a single package to be optimized for all types of use. To this end, *msprime* lacks functionality in ways that limit its utility for studying deeper-scale (e.g., phylogenetic) datasets. Here we describe a new Python package, *ipcoal*, which wraps around *msprime* with the aim of filling this niche: to provide a simple method for simulating genealogies and sequences on species trees or networks.

## 2 Phylogenomic data simulation

We make the following distinctions among terms in *ipcoal*: a genealogy is the true history of ancestry among a set of sampled genes; a gene tree is an empirical estimate of a genealogy based on sequences from some region of the genome; and a species tree is a demographic model including a topology (Maddison, 1997; Pamilo & Nei, 1988). As phylogenetics transitions from a focus on multi-locus data sets (Knowles & Kubatko, 2011) to the analysis of whole genomes – and the spatial distribution of correlated genealogical variation along chromosomes – these distinctions that we highlight in *ipcoal*, between unobservable genealogical variation and the empirical gene tree estimates that can be made from observable sequence data, will become increasingly relevant (Adams & Castoe, 2019; Posada & Crandall, 2002).

Simulating realistic sequence data under the multispecies coalescent model has typically involved a two-step approach: a set of independent genealogies is first simulated, and then a model of sequence evolution is applied along the edges of each tree to produce sequence alignments. This phylogenetic workflow differs from standard population-level coalescent simulations in several ways: (1) phylogenies generally contain many more lineages than population genetic models which makes describing them to coalescent simulators burdensome and error-prone; (2) the phylogenetic workflow typically ignores recombination, but such data can now be simulated easily by modern coalescent software; and (3) the phylogenetic workflow applies a Markov model of sequence evolution rather than the more simple infinite-sites process, allowing for homoplasy and asymmetrical substitution rates. In *ipcoal* we have combined the best aspects of each approach so that it is easy to describe demographic models for large trees, to simulate independent or linked genealogies, and to generate sequences under complex models of sequence evolution.

## 3 Implementation

### 3.1 Reproducible and robust workflow

The *ipcoal* library is designed for interactive use within jupyter-notebooks (Kluyver *et al.*, 2016), where simulations can be run in the same document as downstream statistical analyses; visualization tools can be used to validate model accuracy; and code, figures, and results are easily organized into reproducible and shareable documents. The code is designed to be easy to use, following a minimalist and object-oriented design with few user-facing classes and functions.

### 3.2 Defining demographic models

The primary object that users interact with in *ipcoal* is the `Model` class object (Fig. 1a), which takes a number of user-supplied parameter arguments to initialize demographic and substitution models. The primary convenience of the `Model` class object is its ability to automate the construction of a demographic model by parsing a tree object. For large phylogenies this is important. For example, to describe a demographic model for a species tree with 20 tips in *msprime* would require writing code to define 39 divergence events (MassMigrations). *ipcoal* uses the Python tree manipulation and plotting library *toytree* (Eaton, 2020) to parse, visualize, and annotate trees, making it easy to verify whether variable Ne values and admixture scenarios have been properly defined (Fig. 1a-b).

**Figure 1.**
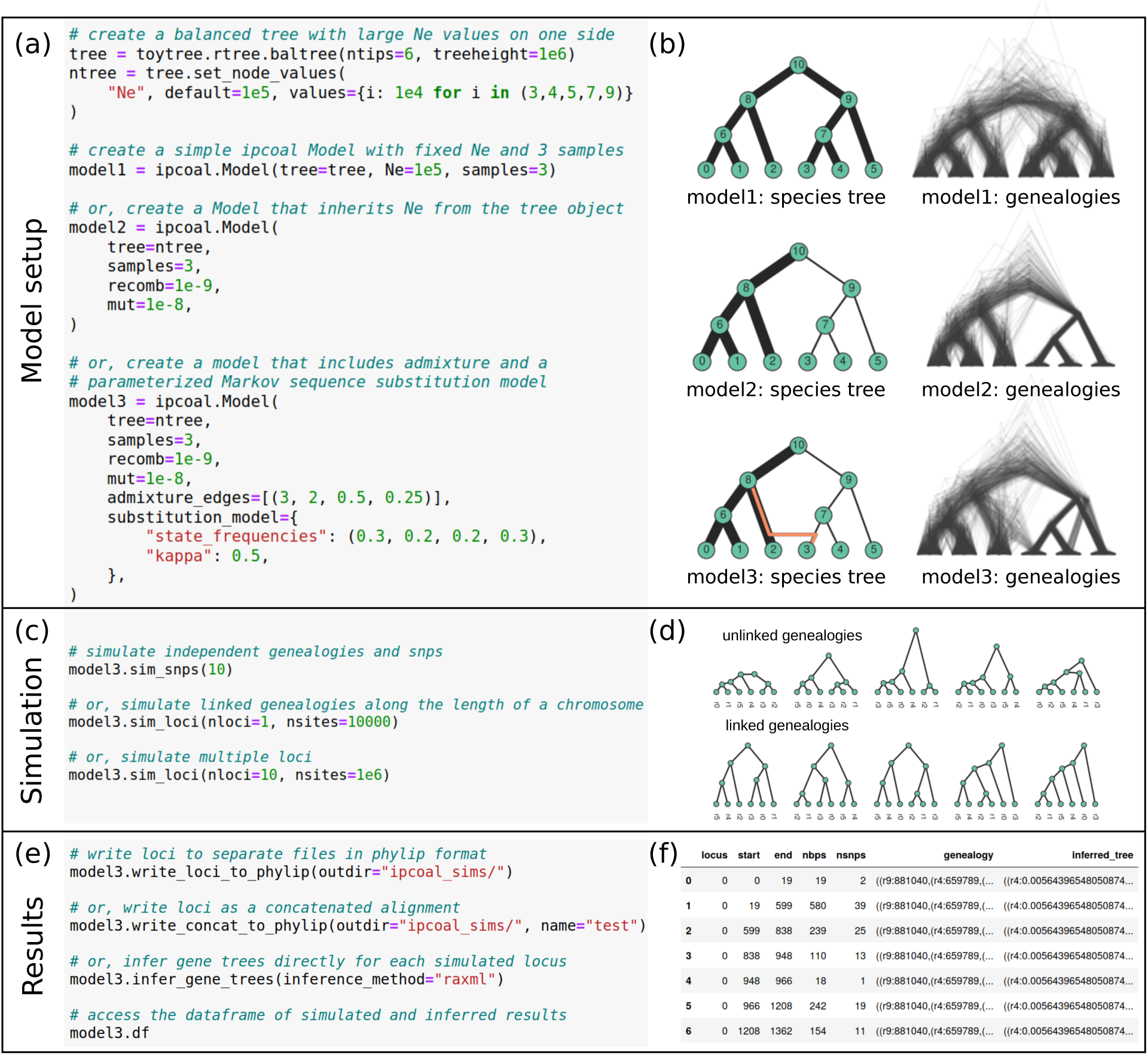
Simulation of coalescent genealogies and sequence data in *ipcoal*. A species tree can be generated or loaded from a newick string to define population relationships in a demographic model, and a single Ne value or variable Nes can be applied to nodes by mapping values using *toytree*. The Model class object of *ipcoal* is used to initialize parameterized demographic and mutational models (a). Genealogical variation reflects parameters of the demographic model including Ne and admixture events, each of which can be easily visualized for validation (b). Sequence data can be simulated as unlinked SNPs (c) or as continuous loci in which recombination affects linkage among neighboring genealogies (d). Simulated sequences can be written to files (either concatenated or as separate loci) for downstream analyses, or the sequences can be used to directly infer gene trees (e). Simulated and inferred results are organized into dataframes for further analyses (f).

### 3.3 Simulating unlinked SNPs

Many inference tools require the input of unlinked single nucleotide polymorphisms (SNPs) to circumvent the effect of recombination (e.g., SVDquartets (Chifman & Kubatko, 2014) and SNAPP (Bryant *et al.*, 2012)). *ipcoal* can generate a distribution of independent genealogies, and unlinked SNPs evolved on those genealogies, using the `Model.sim_snps()` function call (Fig. 1c-d). Notably, we take care that the probability with which a substitution is observed is proportional to the total edge lengths of the genealogy by testing each genealogy for a SNP and moving on to the next independently sampled genealogy if a SNP is not observed. By contrast, users can alternatively toggle the option to enforce a SNP placement on every visited genealogy, which will increase the speed of simulations but introduce a bias toward shallower divergence times.

### 3.4 Simulating loci

The `Model` object can also simulate entire chromosomes (loci) with or without recombination by calling the `Model.sim_loci()` function. This produces sequences of linked genealogies. Nearby genealogies are correlated since some samples share the same ancestors at neighboring genomic regions, and thus are more similar in topology and edge lengths than unlinked trees (Fig. 1d). This type of variation is increasingly of interest for genome-wide analyses.

### 3.5 Simulating sequence evolution

To simulate sequence data on genealogies in *ipcoal*, a continuous-time Markov substitution model is applied iteratively to each edge of the tree from root to tips. We have implemented our own sequence simulator using just-in-time compiled Python code to achieve high performance. We additionally provide the option of using the external tool *seq-sen* (Rambaut & Grass, 1997), which offers a larger range of models than we currently support. Our internal implementation is used by default since it achieves faster speeds by avoiding repeated subprocess calls. The documentation includes test notebooks demonstrating that our implementation converges to the same results as *seq-sen*.

### 3.6 Results

Upon calling a simulation function, two results are stored to the `Model` object: a sequence array (`Model.seqs`) and a dataframe with the genealogy and statistics about each genealogical window (`Model.df`). The sequence array can be written to disk in Nexus or Phylip format, and as separate or concatenated loci, and the DataFrame can be saved as a CSV (Fig. 1e-f). However, to simplify analytical workflows, we provide convenience functions for inferring gene trees directly from sequence data, avoiding the need to organize many files.

## 4 Conclusions

Coalescent simulations for studying genome-wide patterns are routinely used in population genetics, but have not yet achieved widespread use in phylogenetics where the focus has traditionally been limited to a smaller number of unlinked loci. Our new software tool *ipcoal* makes it easy to simulate and explore linked or unlinked genealogical and sequence variation across genomes, providing new opportunities for investigating phylogenetic methods and theory.

## Notes

https://github.com/pmckenz1/ipcoal

https://ipcoal.readthedocs.io/

## References

Adams, R.H. & Castoe, T.A. (2019). Statistical binning leads to profound model violation due to gene tree error incurred by trying to avoid gene tree error. Molecular Phylogenetics and Evolution, 134, 164–171.

Adrion, J.R., Cole, C.B., Dukler, N., Galloway, J.G., Gladstein, A.L., Gower, G., Kyriazis, C.C., Ragsdale, A.P., Tsambos, G., Baumdicker, F., Carlson, J., Cartwright, R.A., Durvasula, A., Kim, B.Y., McKenzie, P., Messer, P.W., Noskova, E., Vecchyo, D.O.D., Racimo, F., Struck, T.J., Gravel, S., Gutenkunst, R.N., Lohmeuller, K.E., Ralph, P.L., Schrider, D.R., Siepel, A., Kelleher, J. & Kern, A.D. (2019). A community-maintained standard library of population genetic models. bioRxiv, p. 2019.12.20.885129.

Beerli, P. & Felsenstein, J. (2001). Maximum likelihood estimation of a migration matrix and effective population sizes in n subpopulations by using a coalescent approach. Proceedings of the National Academy of Sciences, 98, 4563–4568.

Brown, J.M. (2014). Predictive Approaches to Assessing the Fit of Evolutionary Models. Systematic Biology, 63, 289–292.

Bryant, D., Bouckaert, R., Felsenstein, J., Rosenberg, N.A. & RoyChoudhury, A. (2012). Inferring Species Trees Directly from Biallelic Genetic Markers: Bypassing Gene Trees in a Full Coalescent Analysis. Molecular Biology and Evolution, 29, 1917–1932.

Chifman, J. & Kubatko, L. (2014). Quartet Inference from SNP Data Under the Coalescent Model. Bioinformatics, 30, 3317–3324.

Chung, Y. & Hey, J. (2017). Bayesian Analysis of Evolutionary Divergence with Genomic Data under Diverse Demographic Models. Molecular Biology and Evolution, 34, 1517–1528.

Degnan, J.H. & Rosenberg, N.A. (2009). Gene tree discordance, phylogenetic inference and the multi-species coalescent. Trends in Ecology & Evolution, 24, 332–340.

Eaton, D.A.R. (2020). Toytree: A minimalist tree visualization and manipulation library for Python. Methods in Ecology and Evolution, 11, 187–191.

Green, R.E., Krause, J., Briggs, A.W., Maricic, T., Stenzel, U., Kircher, M., Patterson, N., Li, H., Zhai, W., Fritz, M.H.Y., Hansen, N.F., Durand, E.Y., Malaspinas, A.S., Jensen, J.D., Marques-Bonet, T., Alkan, C., Prüfer, K., Meyer, M., Burbano, H.A., Good, J.M., Schultz, R., Aximu-Petri, A., Butthof, A., Höber, B., Höffner, B., Siegemund, M., Weihmann, A., Nusbaum, C., Lander, E.S., Russ, C., Novod, N., Affourtit, J., Egholm, M., Verna, C., Rudan, P., Brajkovic, D., Kucan, Ž., Gušic, I., Doronichev, V.B., Golovanova, L.V., Lalueza-Fox, C., de la Rasilla, M., Fortea, J., Rosas, A., Schmitz, R.W., Johnson, P.L.F., Eichler, E.E., Falush, D., Birney, E., Mullikin, J.C., Slatkin, M., Nielsen, R., Kelso, J., Lachmann, M., Reich, D. & Pääbo, S. (2010). A draft sequence of the neandertal genome. Science, 328, 710–722.

Gronau, I., Hubisz, M.J., Gulko, B., Danko, C.G. & Siepel, A. (2011). Bayesian inference of ancient human demography from individual genome sequences. Nature Genetics, 43, 1031–1035.

Hudson, R.R. (1983). Testing the Constant-Rate Neutral Allele Model with Protein Sequence Data. Evolution, 37, 203–217.

Hudson, R.R. (2002). Generating samples under a Wright–Fisher neutral model of genetic variation. Bioinformatics, 18, 337–338.

Kelleher, J., Etheridge, A.M. & McVean, G. (2016). Efficient Coalescent Simulation and Genealogical Analysis for Large Sample Sizes. PLOS Computational Biology, 12, e1004842.

Kingman, J.F.C. (1982). The coalescent. Stochastic Processes and their Applications, 13, 235–248.

Kluyver, T., Ragan-Kelley, B., Pérez, F., Granger, B.E., Bussonnier, M., Frederic, J., Kelley, K., Hamrick, J.B., Grout, J., Corlay, S., Ivanov, P., Avila, D., Abdalla, S., Willing, C. & al, e. (2016). Jupyter Note-books - a publishing format for reproducible computational workflows. In: ELPUB.

Knowles, L.L. & Kubatko, L.S. (eds.) (2011). Estimating Species Trees: Practical and Theoretical Aspects. 1st edn. Wiley-Blackwell.

Maddison, W.P. (1997). Gene Trees in Species Trees. Systematic Biology, 46, 523–536.

Pamilo, P. & Nei, M. (1988). Relationships between gene trees and species trees. Molecular Biology and Evolution, 5, 568–583.

Posada, D. & Crandall, K.A. (2002). The effect of recombination on the accuracy of phylogeny estimation. Journal of Molecular Evolution, 54, 396–402.

Rambaut, A. & Grass, N.C. (1997). Seq-Gen: an application for the Monte Carlo simulation of DNA sequence evolution along phylogenetic trees. Bioinformatics, 13, 235–238.

Reich, D. (2018). Who we are and how we got here: Ancient DNA and the new science of the human past. Oxford University Press.

Schrider, D.R. & Kern, A.D. (2018). Supervised machine learning for population genetics: A new paradigm. Trends in Genetics, 34, 301 – 312.

